# Allosteric inhibition of PRPS is moderated by filamentous polymerization

**DOI:** 10.1101/2022.04.28.489849

**Authors:** Huan-Huan Hu, Guangming Lu, Chia-Chun Chang, Yilan Li, Jiale Zhong, Chen-Jun Guo, Xian Zhou, Boqi Yin, Tianyi Zhang, Ji-Long Liu

## Abstract

Phosphoribosyl pyrophosphate (PRPP) is an important intermediate for the biosynthesis of purine and pyrimidine nucleotides, histidine, tryptophan, and cofactors NAD and NADP. Abnormal regulation of PRPP synthase (PRPS) has been associated with human disorders including Arts syndrome, retinal dystrophy and gouty arthritis. Recent studies have revealed that PRPS can form filamentous cytoophidia in prokaryotes and eukaryotes. Here we resolve two distinct filament structures of *E. coli* PRPS at near-atomic resolution under Cryo-EM. Formation of two types of filaments is controlled by the binding of different ligands. While the type A filament attenuates the allosteric inhibition of PRPS by ADP, the type B filament enhances the inhibition. In addition, a novel conformation of the regulatory flexible loop of PRPS was found occupying the ATP binding site. AMP/ADP bound at a noncanonical allosteric site interacts with the regulatory flexible loop and facilitates the binding of ATP. Our findings not only reveal molecular mechanisms of the regulation of PRPS with structural basis, but also suggest a distinctive bidirectional regulatory system for PRPP production via PRPS polymerization.

## INTRODUCTION

Phosphoribosyl pyrophosphate (PRPP) is an important intermediate for multiple metabolic pathways in the cell. It is utilized in the biosynthesis of purine and pyrimidine nucleotides, histidine and tryptophan, and the cofactors NAD and NADP^1,2^. PRPP is synthesized by PRPP synthase (PRPS), which catalyzes the transfer of a diphosphate from ATP to ribose-5-phosphate (R5P), thereby generating an AMP and a PRPP^3^.

The proper function of PRPS to maintain the homeostasis of PRPP pool is essential for life. In humans, the missense mutations of PRPP synthase isozyme 1 (PRPS1), which alter the activity or the allosteric regulation of the enzyme, are associated with the cause of many severe pathological outcomes in clinical^4–6^. Among them, the hypoactivity of the enzyme may result in neurological disorders like Arts syndrome, non-syndromic sensorineural deafness 2 (DFN2), Charcot-Marie-Tooth disease 5 (CMTX5), and retinal dystrophy. Conversely, the hyperactivity of PRPS1 could lead to neurosensory defects, hyperuricemia, or gouty arthritis. Misregulated activity and expression of PRPS1 or PRPS2 have been observed in many cancers and correlated with thiopurine drug resistance in relapsed childhood acute lymphoblastic leukemia^7,8^. Thus, the precise regulation of PRPS is of great significance in cell metabolism and physiology.

In general, an organism contains at least one PRPS gene. PRPS in various organisms can be categorized into three classes, according to their distinct biochemical properties. The class I PRPP synthases are the most widely distributed in the phylogeny, from prokaryotes to eukaryotes, including *E. coli* and humans. Phosphate ion (Pi) is required for the activation of class I PRPS, while the ADP, an allosteric inhibitor, competes with Pi and ATP for binding at the allosteric site and the active site, respectively^9–12^. The class II PRPS, however, are active in the absence of Pi. ADP does not bind the allosteric site of the class II PRPS but inhibits it in a competitive manner^13,14^. Class III PRPS, which exists in archaea, requires Pi for activation but has no allosteric mechanism^15^. Therefore, only the class I PRPS is allosterically regulated.

A micron-scale filament of metabolic enzymes, termed the cytoophidium, has been reported in prokaryotes and eukaryotes^16–18^. The cytoophidium is assembled by bundling filamentous polymers of metabolic enzymes. Dozens of cytoophidium-forming enzymes are identified in the genome-wide screening in budding yeast. In mammalian cells, the presence of CTP synthase (CTPS) and IMP dehydrogenase (IMPDH) cytoophidium is proposed to be correlated with distinctive metabolism in specific tissues such as cancers and immune cells. The polymerizations of CTPS and IMPDH have been demonstrated to desensitize the proteins to end-product inhibition or allosteric inhibition effects, suggesting its physiological functions in tuning intracellular nucleotide levels. Recently, PRPS is identified as a novel cytoophidium-forming enzyme in various eukaryotes, including budding yeast, fruit fly, zebrafish, and mammals^19,20^. The evolutionary conservation of PRPS filamentation implies the physiological importance of the structure. Moreover, PRPS serves as the upstream enzyme of CTPS and IMPDH, suggesting that the regulation of enzyme functions via filamentation is particularly important to *de novo* nucleotide biosynthesis. Although high-resolution structures of human, *Bacillus subtilis* and *E. coli* PRPS hexamer have been resolved from a number of crystal forms^12,21–23^, how PRPS assembles the filaments is still obscure. Thus, we aim to investigate the structural basis of PRPS filament and the underlying mechanisms of its functions.

In the current study, we show that *E. coli* PRPS forms filaments *in vitro* and *in vivo*. By using cryo-electron microscopy (cryo-EM), we resolve the structures of PRPS filaments at the near-atomic resolution between 2.3 to 2.9 Å. Structures of two types of PRPS filaments with different ligands are resolved. Structural and biochemical analyses indicate that the formation of type A filaments attenuates the allosteric inhibition, whereas PRPS in type B filaments is more susceptible to the allosteric inhibition. In addition, a noncanonical allosteric binding site for AMP and ADP binding is identified in the type A filament. The binding of AMP/ADP at this site may stabilize the regulatory flexible loop, thereby facilitating the binding of ATP at the active site. Altogether, our findings reveal novel mechanisms for the regulation of *E. coli* PRPS with the structural basis and provide new insights for PRPS-related human disorders and potential clinical and industrial applications.

## RESULTS

### PRPS hexamers assemble two types of filaments

There is only one PRPS gene encoded in the *E. coli* genome, having 47.5% sequence identity to human PRPS1 and PRPS2^24^. Recently, a filamentous macrostructure of PRPS has been observed in various organisms and suggested to be functionally relevant to cell metabolic status. In order to elucidate the functions and underlying molecular mechanisms of PRPS filamentation with a structural basis, we expressed and purified *E. coli* PRPS from *E. coli* K12 strain in Transetta (DE3) cells and performed cryo-EM for the structural analysis.

We first attempted to determine what factors could trigger the filamentation of PRPS. PRPS protein was incubated with various ligands and the filamentation was examined with negative staining. No filament was found when PRPS was incubated without additional ligands. However, while PRPS was incubated with ATP or AMP together with Mg^2+^, many PRPS filaments were observed. In the conditions with only ADP (2 mM), PRPS formed large aggregates of PRPS, which are likely the bundles of PRPS filaments or small crystals. On the other hand, when Pi was mixed with PRPS in different concentrations (10, 30, and 50 mM), filaments were found in samples with 50 mM Pi (Sup 1).

We selected the condition with the supplementation of ATP (2 mM) + Mg^2+^ and the condition with Pi (50mM) to induce PRPS filamentation for structural analysis with cryo-electron microscopy (cryo-EM) and single-particle analysis. As the result, two distinct filament structures, type A and type B filaments were resolved (Figure 1a and b). The type A filaments are assembled in the condition with ATP and Mg^2+^. In this model, hexamers of PRPS pile up in line forming the filamentous polymer. The twist of type A filament is 27° and the rise is 63 Å (Figure 1a). We identify an R5P located at the active site, and a Pi is located at the binding position of the β-phosphate of ADP at the canonical ADP allosteric site (allosteric site 1) (Figure 1a). Surprisingly, instead of an ATP, the active site is occupied by an ADP, and another ADP is located at a binding site (allosteric site 2) that is bound by an AMP in a previous model of *Legionella pneumophila* PRPS (PDB ID: 6NFE).

**Figure 1.**
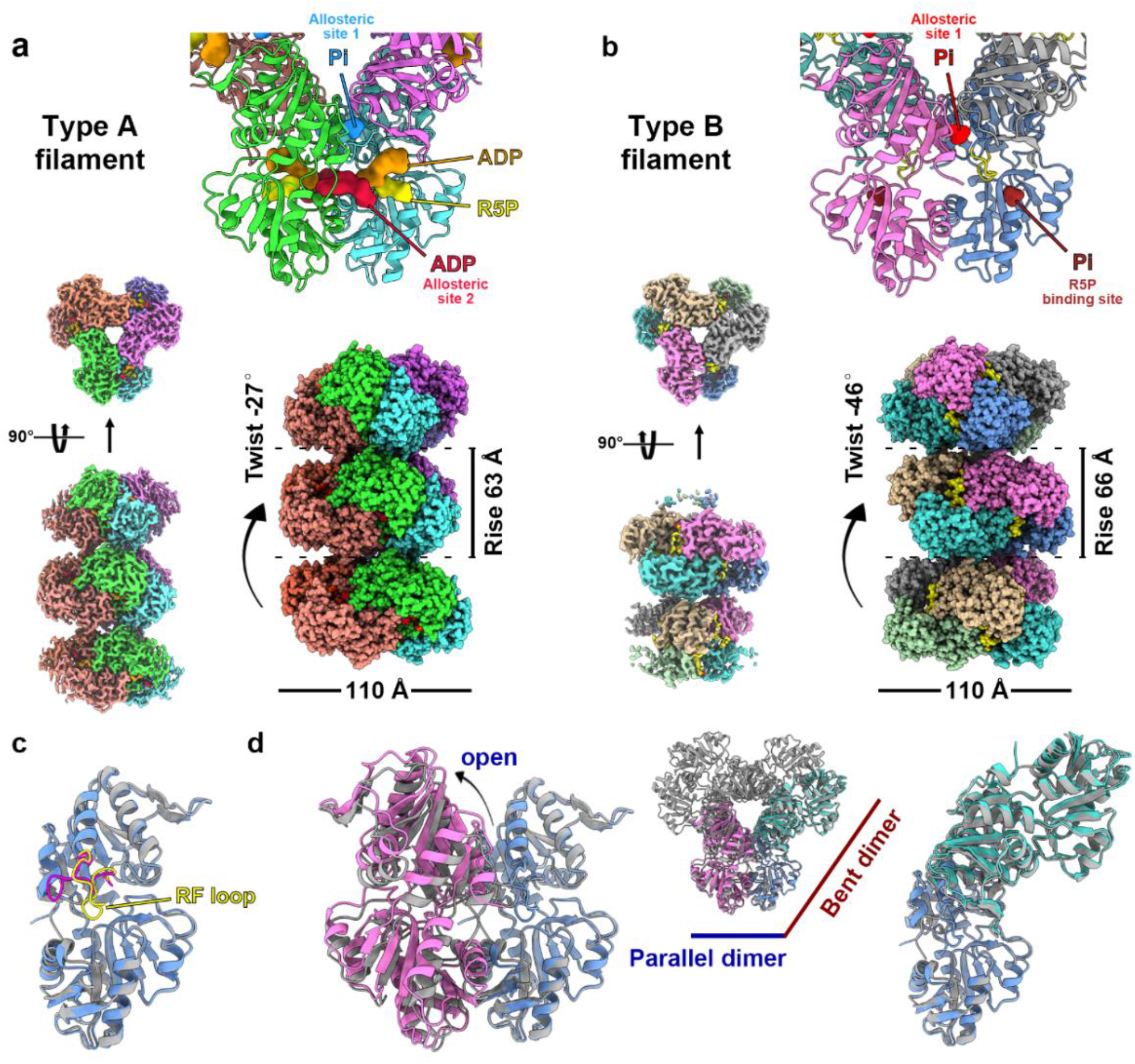
Overall structures of *E. coli* PRPS type A and type B polymers. (a and b) Cryo-EM reconstructions of type A (A, 2.3 Å resolution) and type B (b, 2.9 Å resolution) PRPS polymers. (c) The structural comparison of PRPS monomers in the closed (type A polymer) and open hexamers (type B polymer). The regulatory flexible loop (RF loop) is colored in magenta and yellow in the monomer of type A and type B polymers, respectively (d) The structural comparison of the parallel and bent dimers in the closed (shown in gray) and open hexamers.

Similar to the type A filament, hexamers are the protomers of the type B filament, which is assembled in the condition with Pi (50 mM). The twist of the polymer is 46° and the rise is 66 Å (Figure 1b). The R5P binding site and the allosteric site 1 are bound by Pi, while the ATP binding site and the allosteric site 1 are not bound by any ligand (Figure 1b).

Structural comparison between these two models shows that monomer structures of PRPS in type A and type B filaments are highly similar with an evident conformational change at the regulatory flexible loop (RF loop, Tyr94 to Thr109) (Figure 1c and d). However, the distance between two monomers of the parallel dimer differs in two types of filaments, resulting in a closed hexamer conformation in the type A filament and an open hexamer in the type B filament.

### Ligand binding modes in the type A filament

Being consistent with previous studies, two Mg^2+^ bound in each monomer in type A filament. One Mg^2+^ (Mg site 1) coordinates the hydroxyls of C1, C2, and C3 of R5P with Asp170 and two water molecules. The other Mg^2+^ (Mg site 2) coordinates oxygens of α- and β-phosphates of ADP (active site) with His131 and three water molecules (Figure 2a). The ADP binds to the active site by forming hydrogen bonds with Asp37 and Gln97, a π-π interaction between Phe35 and the adenine base, and salt bridges between Arg99 and His131 and the phosphate. The α-phosphate of the ADP also interacts with R5P through hydrogen bonds with the C-1 hydroxyl. Since we did not add ADP into the mixture for sample preparation, the ADP might be derived from the spontaneous hydrolysis of ATP or preserved during protein purification. A similar scenario may also explain the presence of the R5P in the model, which was not supplemented in any given conditions either.

**Figure 2.**
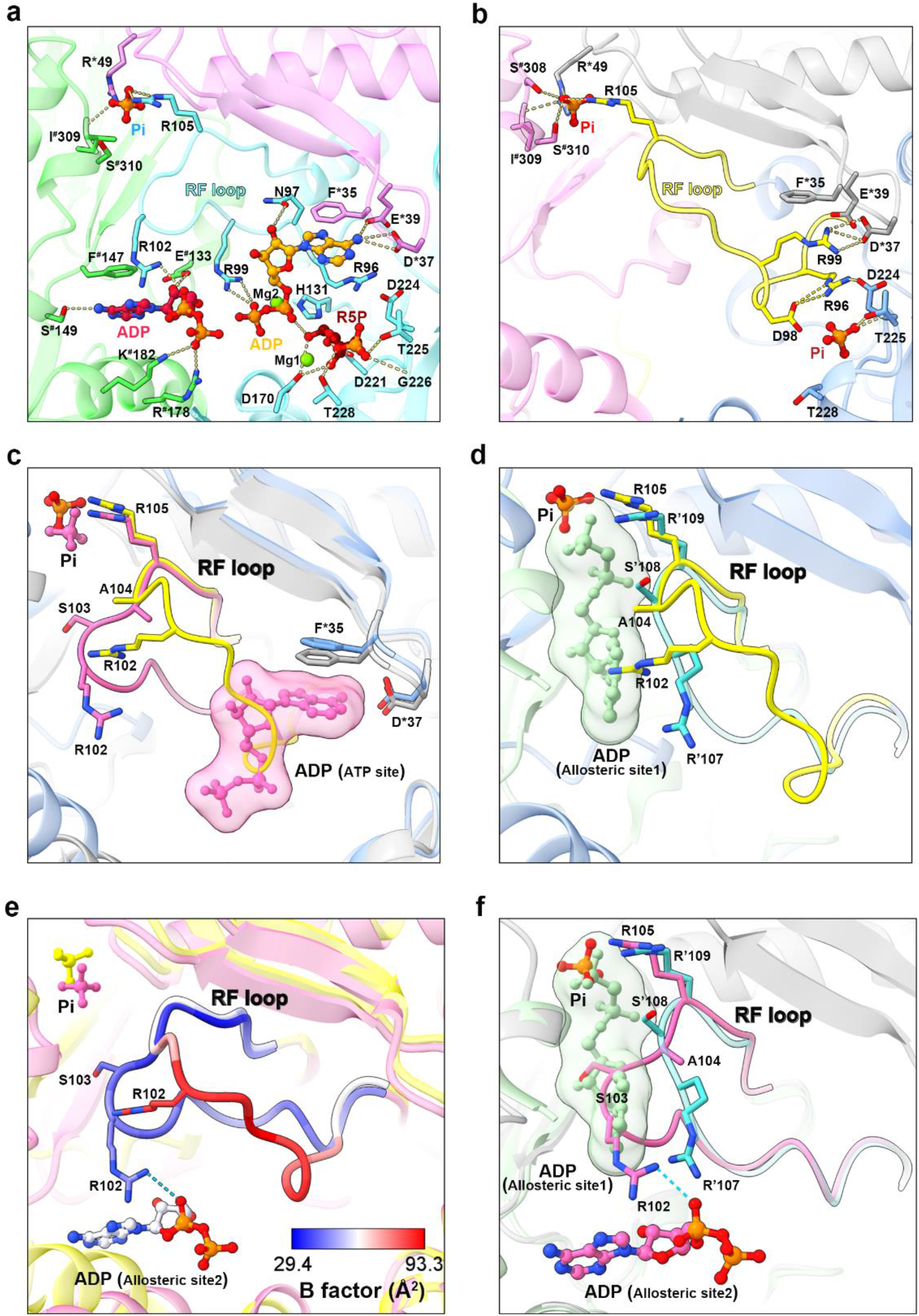
The binding of AMP/ADP at the allosteric site 2 and conformational changes of the regulatory flexible loop imply a novel molecular mechanism of PRPS regulation. (a) ADP and R5P are identified at the active site of PRPS in the type A filament, while the allosteric site is bound by Pi and the allosteric site 2 is bound by an ADP. Residues interacting with ligands are indicated. (b) The ATP binding site is not bound by any ligand, while the R5P binding site and the allosteric site 1 are bound by Pi in the type B filament. (c) The structural comparison of the RF loop in type A (pink and gray) and type B (yellow and blue) filaments. The RF loop of the type B filament overlaps with the ADP at the active site in the type A filament. (d) The structural comparison of the RF loop in the type B filament and the *Bacillus substilis* PRPS (PDB ID: 1DKU). (e) B factors are shown on the RF loop in the models of type A and type B filaments. (f) The structural comparison shows the RF loop in the type A filament overlaps with the ADP at the allosteric site 1 in the model the *Bacillus substilis* PRPS (PDB ID: 1DKU).

The allosteric site 2 is not bound by any ligands in previous bacterial PRPS models except a *Legionella pneumophila* PRPS model (PDB ID: 6NFE), in which the binding site is bound by an AMP in the same fashion, suggesting this binding site can accommodate both AMP and ADP. We resolved another type A filament model (type A^AMP/ADP^) with PRPS pre-incubated in the condition supplemented with AMP (2 mM) and ADP (2 mM) (Sup 2). This structure is nearly identical to the former type A filament model with an AMP at the allosteric site 2 and an empty allosteric site 1 (Sup 2). The ADP binds the allosteric site 2 by forming hydrogen bonds with Arg102, Ser149, and Glu133, a π-π interaction between Phe147 and the guanine base, and salt bridges between Arg102, Arg178, Lys182, and the phosphates. (Sup 2, Figure 2a). In some human PRPS1 models, the allosteric site 2 is bound by an SO_4_^2-^ and residues responsible for the binding of AMP/ADP are conserved between *E. coli* and human PRPS isoforms, implying the potential evolutionary conservation of this allosteric binding site (Sup 3, sequence comparison).

### Conformational changes of the regulatory flexible loop reveal novel regulatory mechanisms of PRPS

In the model of type B filament, the allosteric site is bound by a Pi and another Pi is located at the binding position for the phosphate of R5P, while the ATP binding site is not bound by any ligand (Figure 2b). Intriguingly, our density map shows that the ATP binding site is occupied by the RF loop (Figure 2c and Sup 4). In the model of *Bacillus substilis* PRPS with ADP at the allosteric site 1 (PDB ID: 1DKU), the RF loop is not present at the same position as that in the type B filament (Figure 2d). Thus, this conformation may suggest a novel role of the RF loop in the regulation of ATP binding.

The RF loop is unstable in the type B filament with the B-factor ranging from -50.5 to -93.3 Å^2^ (Figure 2e). However, it is stabilized by multiple interactions including the salt bridge between Arg102 and the ADP at allosteric site 2 with the B-factor ranging from -29.4 to -44.5 Å^2^ in the type A filament (Figure 2f). In the stable conformation, the RF loop is anchored at the allosteric site 1 and covers the ADP binding pocket, resulting in a closed structure unavailable for ADP binding (Figure 2f). Therefore, we propose that the conformational changes of the RF loop may also control the binding of ADP at the allosteric site 1. This may explain the fact that the allosteric site 1 in the type A^AMP/ADP^ filament model is not bound by ADP even though its binding competitor Pi is not present at the binding site (Sup 2).

### Distinct contact sites of PRPS hexamers in type A and type B filaments

Hexamers in the type A filament are connected by the salt bridges formed between pairs of Arg301 and Glu298, the hydrogen bonds between Arg301, Asn305, and Glu307 of two neighboring hexamers, and also by the van der Waals’ Forces between Arg302 and Arg301 of two hexamers (Figure 3a). In contrast, hexamers in the type B filament polymerize through the π-cation interaction between Tyr24 and Arg22, and the hydrogen bond between Arg301 and Leu23 of neighboring hexamers. (Figure 3b). Notably, it is unlikely that a single filament could accommodate both open and closed hexamers since the pairing of two hexamers depends on distinct contact sites in these two conformations. In order to investigate the functions of both types of filaments, we generated PRPS^R302A^ and PRPS^Y24A^ to disrupt the formation of type A and type B filaments, respectively. The mutant PRPS carrying both R302A and Y24A mutations (PRPS^R302A/Y24A^), which is supposedly unable to form either type of filament was also generated. The capabilities of each PRPS mutant for forming type A and type B filaments were assessed with negative staining. To our expectation, PRPS^R302A/Y24A^ failed to assemble filaments in all conditions, and PRPS^R302A^ and PRPS^Y24A^ can only form type B and type A filaments, respectively. (Figure 3c).

**Figure 3.**
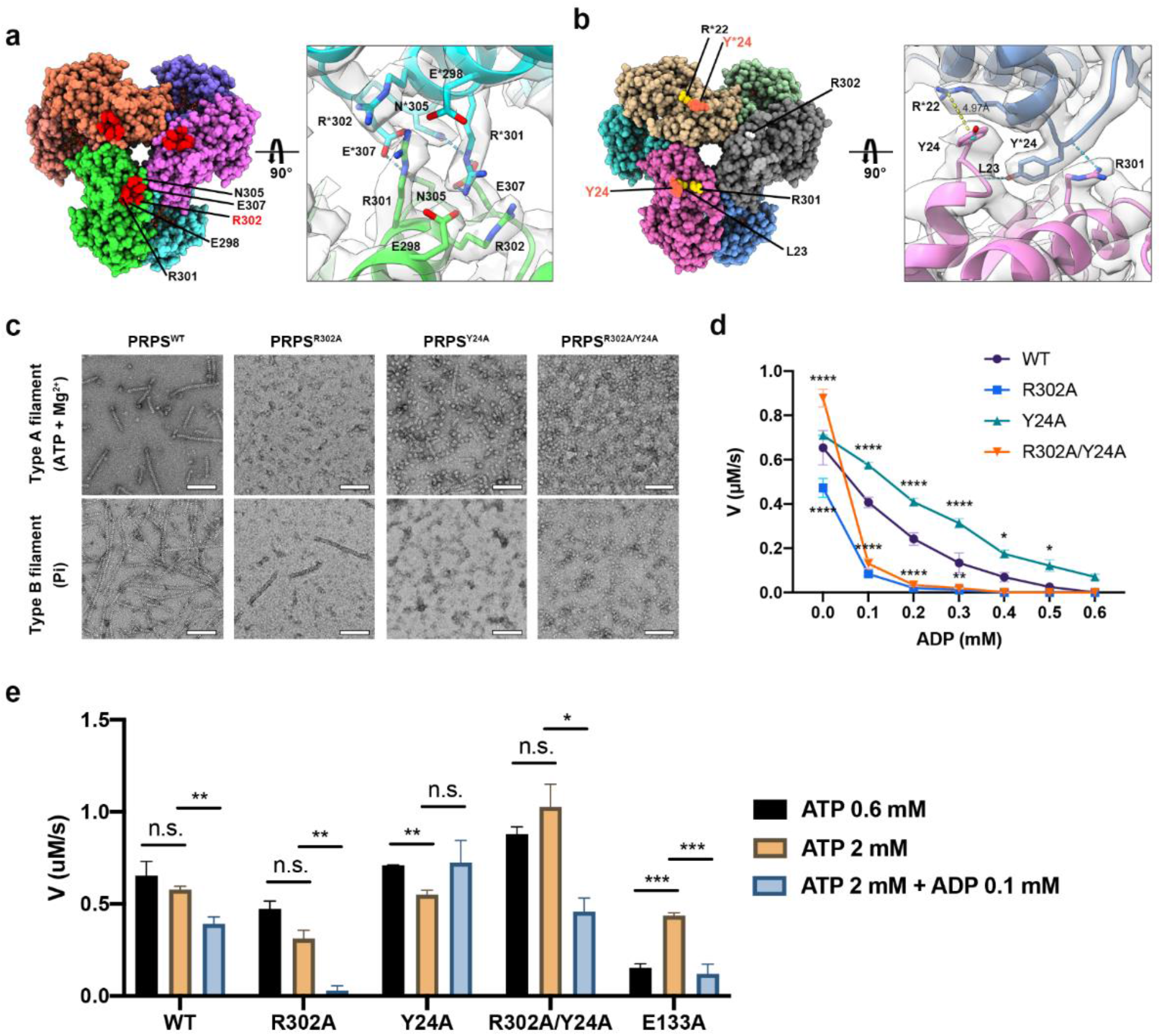
The allosteric inhibition of *E. coli* PRPS is regulated by filamentation. (a and b) Maps and models of type A (a) and type B (b) filaments reveal distinct interfaces for the interaction between neighboring hexamers in two types of filaments. Residues responsible for the interactions are indicated. (c) Negative staining images of wildtype and mutant *E. coli* PRPS in the conditions that induce type A and type B filaments. Scale bars = 100 nm. (d) The graph shows the catalytic activity of wild-type and mutant *E. coli* PRPS in conditions with various amounts of ADP. (Tukey’s test) (e) A bar graph shows the catalytic activity of wild-type and mutant *E. coli* PRPS in the reaction mixtures with different amounts of ATP and ADP. (Student’s *t-test*) Error bars = S.E.M.

### Filamentation regulates the allosteric inhibition of PRPS

In order to elucidate whether disruption of PRPS filamentation would affect its catalytic activity and allosteric regulation, we measured the in vitro PRPP production with a coupled reaction. In the reaction mixture, newly synthesized PRPP for the reaction catalyzed by PRPS would be subsequently utilized by phosphoribosyltransferase (OPRT) in the reaction OA + PRPP → OMP + PPi^25^. Thus, the PRPP production can be determined by the consumption of OA, which has an absorbance at 295 nm. In the conditions with no ADP, the activity of PRPS^WT^ shows no significant changes in concentrations of ATP and R5P higher than 0.6 and 0.5 mM (Sup 5), respectively. It is known that the catalysis of class I PRPS requires Pi as an activator.

In the absence of Pi, the activity is undetectable. When Pi was added into the reaction mixtures at different levels, the activity peaked at 5 mM and gradually decreased as the concentration of Pi increased (Sup 5). Therefore, we used 0.6 mM ATP, 0.6 mM R5P, and 10 mM Pi in combination with different concentrations of ADP to analyze the effects of individual mutations in the activity and the allosteric inhibition.

The activity of wildtype and each mutant PRPS was measured in the conditions with ADP in a concentration gradient. When there was no ADP, PRPS^R302A^ displayed significantly lower activity in comparison with PRPS^WT^, whereas PRPS^R302A/Y24R^ showed higher activity (Figure 3d). When ADP (0.1 mM) was added into the mixture, the activity of PRPS^R302A^ and PRPS^R302A/Y24A^ dramatically dropped by 82.3% and 85.1%, whereas the activity of PRPS^WT^ and PRPS^Y24A^ dropped by only 37.8% and 18.9%, respectively (Figure 3d). Along with increased ADP concentrations, both PRPS^WT^ and PRPS^Y24A^ were gradually inhibited and the activity of PRPS^R302A^and PRPS^R302A/Y24R^ was nearly undetectable when ADP concentration was higher than 0.2 mM.

ADP is known to inhibit PRPS through both allosteric and competitive inhibition. The inhibitory effect we observed in given conditions should be a combination of both mechanisms. For the sake of reducing the competitive inhibition, we increased ATP concentration to 2 mM (Figure 3e). The higher ATP concentration did not change the activity of all PRPS when ADP was not present. However, when ADP (0.1 mM) was supplied, nearly all groups displayed a reduction of activity (Figure 3e). The activity of PRPS^WT^ and PRPS^R302A^ dropped by 32.2% and 91.6%, respectively. The only exception is PRPS^Y24A^, of which the activity was unchanged. The activity of PRPS^R302A/Y24A^ also had a 55.4% reduction when ADP was added, indicating that the formation of type B filament is not required for allosteric inhibition. These results show that the disruption of type A filament enhances the allosteric inhibition, while the loss of type B filament leads to remarkable resistance to allosteric inhibition. Thus, we propose that the type A filament structure functions to maintain PRPS in the closed conformation, in which the allosteric site 1 is not accessible for ADP, thereby desensitizing the allosteric inhibition. By contrast, PRPS hexamers in type B filaments are stabilized in the open conformation, which might be more susceptible to ADP binding at the allosteric site 1, and consequently enhance the inhibitory effect.

### AMP/ADP at the allosteric site 2 may facilitate ATP binding at the active site

One of the major structural differences between our two models is the integrity of the allosteric site 2. In order to further address the significance of the binding of AMP/ADP at the allosteric site 2, we replaced ADP in the reaction mixtures with AMP and found no significant change in the activity in the conditions with different AMP levels (Sup 5d). Since AMP is supposedly unable to bind at the allosteric site 1 and the active site, our results indicate that the binding of AMP/ADP at the allosteric site 2 does not induce catalytic inhibition.

According to our models, this AMP/ADP at allosteric site 2 may participate in the stabilization of the RF loop. Since the stabilization of the RF loop may expose the ATP binding site, loss of the AMP/ADP at allosteric site 2 is expected to attenuate ATP binding affinity. We intended to impair the interaction between hydroxyls of C2 of ADP and the side chain of Glu133 by introducing a point mutation, E133A. When ATP was supplied in 0.6 mM, the activity of PRPS^E133A^ was four times lower than that of PRPS^WT^. However, when ATP in the reaction mixtures was increased to 2 mM, the activity of PRPS^E133A^ was restored to a comparable level to PRPS^WT^, suggesting that the reduced AMP/ADP binding affinity of the allosteric site 2 results in a decrease in the ATP binding affinity of the active site (Figure 3e).

### *E. coli* PRPS filaments assemble the cytoophidium in vivo

Filamentous polymers of various metabolic enzymes have been shown to bundle into a large filamentous structure, which is denoted the cytoophidium, in a broad spectrum of organisms. PRPS cytoophidia have already been observed in multiple eukaryotes including mammals. We fused the sequences of PRPS^WT^ and PRPS^R302A^, PRPS^Y24A^ and PRPS^R302A/Y24A^ mutants were with mCherry and overexpressed them in *E. coli* individually. In a small subset of cells (less than 1%), filamentous structures of PRPS^WT^-mCherry could be observed (Figure 4a). In contrast, PRPS^R302A^-mCherry filaments and dot-like aggregates were observed in some cells. In cells expressing PRPS^Y24A^-mCherry, no filaments but dot-like aggregates were found in some cells in all stages of growth, while none of filaments or dots were observed in cells expressing PRPS^R302A/Y24A^-mCherry. Notably, the width and length of PRPS polymer are within a range of dozens of nanometers, the filaments or dots we observed with fluorescence microscopy are presumably structures on a larger scale. The dots could also comprise PRPS polymers and the polymers may possibly present in cells with no detectable aggregates.

**Figure 4.**
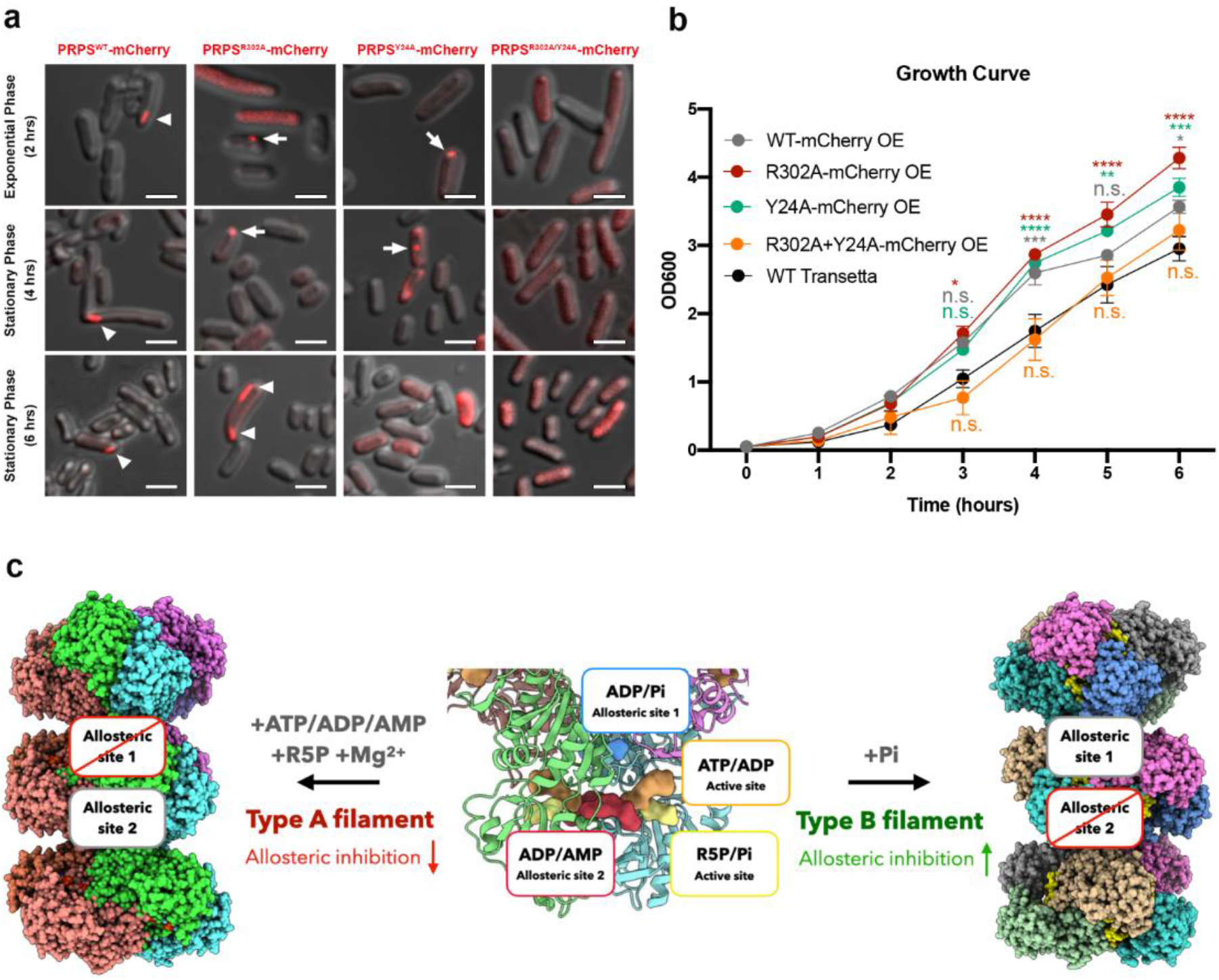
*E. coli* PRPS forms type A and type B filaments in vitro and in vivo. (a) Representative images of Transetta *E. coli* strains overexpressing wildtype and mutant PRPS-mCherry fusion proteins. The filamentous cytoophidia and dot-like aggregates were indicated by arrowheads and arrows, respectively. (b) Growth curve of wildtype Transetta cells and cells overexpressing wildtype and mutant PRPS-mCherry fusion proteins. (Tukey’s test) Error bars = S.E.M. (c) Schematic model of the regulation and function of *E. coli* PRPS filaments.

We next analyzed the growth curve of *E. coli* overexpressing PRPS-mCherry proteins. Cells that overexpress PRPS^WT^, PRPS^R302A^, and PRPS^Y24A^ grew significantly faster than wildtype Transetta cells, while cells overexpressing PRPS^R302A/Y24A^ displayed a similar growth rate to wild-type cells (Figure 4b). Although we could not avoid the possible effects from the overexpression of PRPS and the mCherry-tag, which could potentially influence the formation of the cytoophidium and cell growth, our findings suggest that both type A and type B PRPS filaments are present in vivo and may participate in the coordination between cell metabolism and cell growth.

## DISCUSSION

Assembly of filamentous polymers has emerged as a common and conserved regulatory mechanism for many metabolic enzymes in eukaryotes, prokaryotes, and even archaea^16–18^. Herein we demonstrate two types of filament structures of *E. coli* PRPS. The hexameric propeller structure of PRPS and the residues identified responsible for the polymerization are conserved between *E. coli* and human PRPS1/2, suggesting the property of forming type A and/or type B PRPS filaments is also present in mammalian models. In fact, the isolation of PRPS from rat liver tissues revealed a heterogenous protein complex with a molecular mass larger than 1000 kDa^26^. A similar phenomenon was seen in human PRPS1/2 isolation from tissue sources as well^27,28^. Furthermore, human PRPS1/2 recombinants purified from *E. coli* can also spontaneously assemble into large complexes in vitro with a size larger than 1000 kDa^10^. Although it is not necessarily that these complexes are substantially identical to the filaments depicted in this study, such pieces of evidence together with our findings suggest that class I PRPS is widely under the regulation of the assembly of large complexes in bacterial and mammalian organisms.

Although some filamentous polymers of metabolic enzymes were demonstrated to accommodate different states of the proteins, their protomers in various states assemble through the same interfaces and can either boost or suppress the activity^29–32^. With structural and biochemical approaches, we reveal a distinctive regulatory machinery of E. coli PRPS polymerization. In our findings, two distinct PRPS filament types display opposite directions in the regulation of PRPS activity (Figure 3 and 4c). PRPS hexamers in type A and type B filament are connected through different contact interfaces. It is presumed that one filament could only accommodate hexamers in either closed or open conformation. In well-defined in vitro environments, it is evident that only type A or type B filament could be formed in one particular condition. However, it is plausible that both types of filaments may co-exist in vivo. Our model suggests a more complicate system of protein filamentation in E. coli PRPS and this may contribute to the precision in fine-tuning the production of PRPP, which is the central metabolite in multiple metabolic pathways.

The determinant of forming either open or closed hexamers is proposed to be depending on the ligands at two allosteric sites as the binding of ADP at the allosteric site 1 and AMP/ADP at the allosteric site 2 could possibly stabilize the hexamer in the open and closed state, respectively. Once either allosteric site is bound by an adenine nucleotide, the other will be no longer available for the binding of nucleotides. The interactions between two neighboring hexamers in the filament may also contribute to the maintenance of a particular conformation. Indeed, it has been demonstrated by equilibrium binding dialysis that ATP binds to the active site in the presence of R5P. The same site could be bound by ADP regardless R5P is present or not, and ADP binds to the additional allosteric binding site to yield two bound ADPs per monomer in the presence of R5P^33^. According to our models, both allosteric site 1 and site 2 could be responsible for the binding of the second ADP in these conditions.

In a comparison of these two models, we identified the conformational changes of the RF loop. The loop covers the allosteric site 1 in the closed hexamers of the type A filament. In the open hexamers of the type B filaments, the loop structure is flexible and located at the ATP binding site. In type A filaments, the allosteric site 2 is demonstrated to accommodate both AMP and ADP. The interaction between the bound AMP/ADP and the RF loop suggests the binding of substrate ATP is regulated by the allosteric site 2. This notion could be supported by our activity analysis with the mutant PRPS^E133A^, which displayed a significantly reduced activity when a lower ATP concentration (0.6 mM) was applied to the reaction.

ADP has been known as an effective inhibitor of PRPS as it downregulates the activity of rat PRPS by roughly 40∼50% in a concentration of 0.1 mM. In agreement with previous studies, we show that *E. coli* PRPS^WT^ activity dropped by about 40% when 0.1 mM ADP was added into the reaction mixture containing saturated ATP^9^. We show that such reduction in the activity is compromised by the formation of type A filaments. The disruption of type A filament by the point mutation R302A resulted in nearly undetectable activity in the same condition. Since the intracellular ADP level is generally within the range of hundreds micromolar in *E. coli* and mammalian cells, it is possible that assembly of type A filament is essential for efficient production of PRPP in the cell^34^.

In contrast, type B filament is demonstrated to enhance the allosteric inhibition. The point mutation, Y24A, which disrupts the interface between hexamer in the type B filament, strikingly desensitizes *E. coli* PRPS to the allosteric inhibition by ADP binding. We show that PRPS^Y24A^ retains the full activity in the condition with 0.1 mM ADP and saturated ATP. PRPS mutants have been employed for the improvement of the production of commercial compounds for which PRPP is an intermediate. For instance, feedback-resistant mutant PRPS has been shown to increase the synthesis of riboflavin and purine nucleosides by *R. gossypii* and *B. amyloliquefaciens*, respectively^35,36^. Thus, the point mutations that enhance the formation of type A filament or that prevent the formation of type B filament are potentially applicable to the industry.

In humans, mutations in PRPS1 that render hypoactive or hyperactive of PRPP synthesis have been associated with various disorders. Some of these human PRPS1 mutants have been characterized at the molecular level^4^. The kinetics of inhibition indicated that they were less sensitive to the allosteric inhibition. Interestingly, mutations that lead to desensitization to allosteric inhibition, such as D51H, L128I, D182H, A189V, and H192D, are almost exclusively in the interface between dimers in the hexamer^4^. It is reasonable to speculate that the alternation of the propensity for forming closed or open hexamer conformation may partially explain the molecular mechanisms of particular PRPS variants.

The filamentous macrostructure cytoophidium has been regarded as large bundles of filamentous polymers of metabolic enzymes^16,17^. The presence of various types of cytoophidia has been proposed to be correlated with the specific cellular status in certain tissues. For instance, CTPS cytoophidia are widely displayed by *Drosophila* tissues, especially in proliferative cell types^37–39^. In mammalian models, CTPS cytoophidia were found in mouse thymus and multiple human cancers^40,41^. The cytoophidium may also serve to protect its component proteins from degradation^42,43^. Although the structures of CTPS filaments have been resolved, and the ligands required for CTPS polymerization are determined, the mechanisms of the regulation of CTPS filaments bundling are not well understood^29,31,44,45^. Many factors including the mTOR pathway, temperature, pH value, osmolality, and protein post-translational modifications have been demonstrated to influence the assembly of CTPS cytoophidia in various organisms^31,43,46–50^. Recently, PRPS cytoophidia have been reported in yeast, *Drosophila*, zebrafish, and mammalian cell lines^19,20^. Likewise, we show that PRPS cytoophidia assembled in *E. coli* in different growth phases. The assembly of the cytoophidium reflects the fact that many PRPS filaments are present in the cell. Apart from regulating the allosteric inhibition, whether the bundling of filaments can modulate other properties of PRPS requires further investigation.

In sum, our findings not only reveal that filamentation of PRPS plays a vital role in the cell metabolism, but also suggest a novel mechanism of the regulation of ATP binding by the RF loop. These results expand our understanding of the regulation of the key step in nucleotide biosynthesis and shed light on potential applications in the clinic and industry.

## STAR METHODS TEXT

### *E. coli* PRPS Expression and Purification

Full-length of wildtype or mutant *E. coli* PRPS sequences with a C-terminal 6×His-tag were cloned into a modified pRSFDuet vector at the MCS 2 site and expressed in *E. coli* Transetta (DE3) cells. Cells were cultured at 37 °C for 4 hours after induction with 0.1 mM IPTG at the OD600 range of 0.5∼0.8 and pelleted by centrifugation at 4,000 r.p.m. for 20 minutes. All the remaining purification procedures were performed at 4 °C. The harvested cells were resuspended in cold lysis buffer (50 mM Tris HCl pH8.0, 500 mM NaCl, 10% glycerol, 10 mM imidazole, 5 mM β-mercaptoethanol, 1 mM PMSF, 5 mM benzamidine, 2 μg/ml leupeptin and 2 μg/ml pepstatin). The cell lysate was then centrifuged (18,000 r.p.m.) at 4 °C for 45 minutes after ultrasonication. The supernatant was collected and incubated with equilibrated Ni-NTA agarose beads (Qiagen) for 1 hour. Subsequently, the column was further washed with lysis buffer supplemented with 50 mM imidazole. Target proteins were eluted with elution buffer containing 50 mM Tris HCl pH8.0, 500 mM NaCl, 250 mM imidazole, and 5 mM β-mercaptoethanol. Further purification was performed by using HiLoad Superdex 200 gel-filtration chromatography (GE Healthcare) in column buffer (25 mM Tris HCl pH 8.0 and 150 mM NaCl). The peak fractions were collected, concentrated, and stored in small aliquots at −80 °C.

### Cryo-EM Grid Preparation and Data Collection

To generate type A filaments, 6 μM PRPS protein was dissolved in the buffer containing 25 mM Tris HCl pH 7.5, 2 mM ATP, 10 mM MgCl2. The ATP was replaced with 2 mM ADP and 2 mM AMP for generating type A^AMP/ADP^ filaments. For type B filaments formation, 6 μM PRPS protein was incubated in the buffer containing 50 mM Na2HPO4 and 100 mM NaCl. All the samples were incubated at 37 °C for 30 minutes before being loaded on grids. For the preparation of cryo-EM grids, protein samples were loaded on fresh glow-discharged 300-mesh amorphous alloy grids (CryoMatrix M024-Au300-R12/13). Grids were blotted for 3.5 sec with blot force of -1 at 4 °C and 100% humidity before plunge-freezing in liquid ethane with an FEI Vitrobot Mark IV (ThermoFisher Scientific).

Micrographs were collected at 300 kV on an FEI Titan Krios electron microscope with a K3 Summit direct electron camera (Gatan) in super-resolution counting mode. Automated data acquisition was performed with SerialEM^51^ at a nominal magnification of 22,500×, corresponding to a calibrated pixel size of 1.06 Å with a defocus range from 1.0 to 2.5 μm. Each movie stack was acquired in a total dose of 60 *e*^−^Å^−2^, subdivided into 50 frames at 4 s exposure.

### Image Processing

All image processing steps were performed using Relion3.1-beta^52^. Beam-induced motion correction and exposure weighting were performed by the MotionCorr2^53^ and the CTF (contrast transfer function) parameter was estimated by CTFFIND4. For the dataset of the type A filament, 3045 images were selected manually and 887654 particles were auto-picked. Among them, 438771 particles were selected for 3D classification after 2 rounds of fast 2D classification with particle extracted at binning 2 and another round of 2D classification for binning 1. The featureless cylinder was reconstructed using the relion_helix_toolbox command and applied as a reference model for 3D classification. After 2 rounds of 3D classification with C1 and D3 symmetry applied, a total of 70541 particles of the best class were selected for 3D auto-refinement, and CTF refinement and Bayesian polishing were performed on each particle, resulting in an initial 2.8 Å density map include 3 layers of PRPS hexamer. A final 2.3 Å map was sharpened by post-process using a tight mask for the central hexamer with a B-factor of -45 Å^2^.

For the type B filament dataset, a similar procedure was performed. A total of 1186879 particles were auto-picked from 2776 images. After multiple rounds of 2D classification and 3D classification,168218 particles were selected for the 3D auto-refinement, and CTF refinement and Bayesian polishing were performed for each particle. The best density map was sharpened to a nominal resolution of 2.9 Å by using a tight mask for the central hexamer with a B-factor of -98 Å^2^.

For the type A^AMP/ADP^ filament dataset, a total of 1066797 particles were auto-picked from 1824 images. After multiple rounds of 2D classification and 3D classification,53045 particles were selected for the 3D auto-refinement, and CTF refinement and Bayesian polishing were performed for each particle. The best density map was sharpened to a nominal resolution of 2.6 Å by using a tight mask for the central hexamer with a B-factor of -51 Å^2^. LocalRes was used to estimate the local resolution for all the maps.

### Model Building and Refinement

The Crystal structure of *E. coli* PRPS [Protein Data Bank (PDB) ID: 4S2U] was applied for the initial models of all datasets. The hexamer models were separated and docked into the corresponding electron density map using Chimera v.1.14^54^, followed by iterative manual adjustment and rebuilding in Coot^55^ and real-space refinement in PHENIX^56^. The final atomic model was evaluated using MolProbity^57^. The map reconstruction and model refinement statistics are listed in Supplementary Table 1. All the figures and videos were generated using UCSF Chimera and ChimeraX^58^.

### PRPS Activity Assays

The activity of PRPS was measured with a coupled continuous spectrophotometric assay in a 96-well plate using SpectraMax i3. The PRPS reaction (ATP + R5P → PRPP + AMP) was coupled with *E. coli* orotate phosphoribosyltransferase (OPRT, EC 2.4.2.10) forward reaction (OA + PRPP → OMP + PPi) and the amount of PRPP generated in the reaction was determined by the decrease in orotate (OA) in the mixture. The concentration of OA was measured by the absorbance at 295 nm for 300 sec at 25 °C^25^. The reaction mixture (200 μl) contains 0.1 μM PRPS, 10 mM OPRT, 1 mM OA, 10 mM MgCl2, 1 mM orotate, 250 mM NaCl, 10 mM Na2HPO4, 0.6 mM R5P and AMP, ADP, ATP in concentrations as described in each experiment. ATP or R5P was least added into the mixture to initiate the reaction. All measurements were performed in triplicate.

### Negative staining

Purified *E. coli* PRPS protein (1 μM) was dissolved in Tris-HCl buffer (25 mM Tris-HCl, 150 mM NaCl, 10 mM MgCl2, and 2 mM ATP or ADP) or Pi buffer (50 mM Na2HPO4,300 mM NaCl). After 1-hour of incubation at 37°C, the protein sample was loaded onto hydrophilic carbon-coated grids and washed twice with uranium formate. Subsequently, the grids were dyed with uranium formate. Imaging was acquired with a 120 kV electron microscope (Talos L120C, ThermoFisher, USA) with an Eagle 4 K × 4 K CCD camera system (Ceta CMOS, ThermoFisher, USA) at 57,000 × magnification.

### Sample preparation and confocal microscopy

*E. coli* cells were fixed with 4% formaldehyde for 10 minutes at 37°C, 220 r.p.m. After fixation, cells were collected by centrifugation for 1 minute at, 12000 r.p.m. Cell pellets were washed with PBS twice and resuspended in PBS containing Hoechst33342. Cells were then incubated at room temperature for 1 hour. 2.5 μL of cell solution was mixed with 1 μL of pre-melted 1.2% low melting-point agar on a glass slide and covered with a coverslip for observation. Images were captured under Plan-Apochromat 63×/1.40 Oil DIC M27 objective on a Carl Zeiss LSM 800 (Axio Observer Z1) inverted fluorescence confocal microscope.

### Growth curve

*E. coli* cells were pre-cultured at 37°C, 220 r.p.m. in 2 mL LB medium overnight, and then inoculated at OD600 = 0.05 in a 5 mL LB culture at 37°C, 220 r.p.m.

The growth of the cell is determined by the value of OD600, which is measured by Eppendorf BioPhotometer D30 every hour after inoculation.

## ACKNOWLEDGEMENTS

The EM data were collected at the ShanghaiTech Cryo-EM Imaging Facility. We thank the Molecular and Cell Biology Core Facility (MCBCF) at the School of Life Science and Technology, ShanghaiTech University for providing technical support. This work was supported by grants from Ministry of Science and Technology of China (No. 2021YFA0804701-4), National Natural Science Foundation of China (No. 31771490), Shanghai Science and Technology Commission (No. 20JC1410500) and the UK Medical Research Council (grant nos. MC_UU_12021/3 and MC_U137788471) for grants to J.L.L.

## SUPPLEMENTARY FIGURES

**Figure S1.**
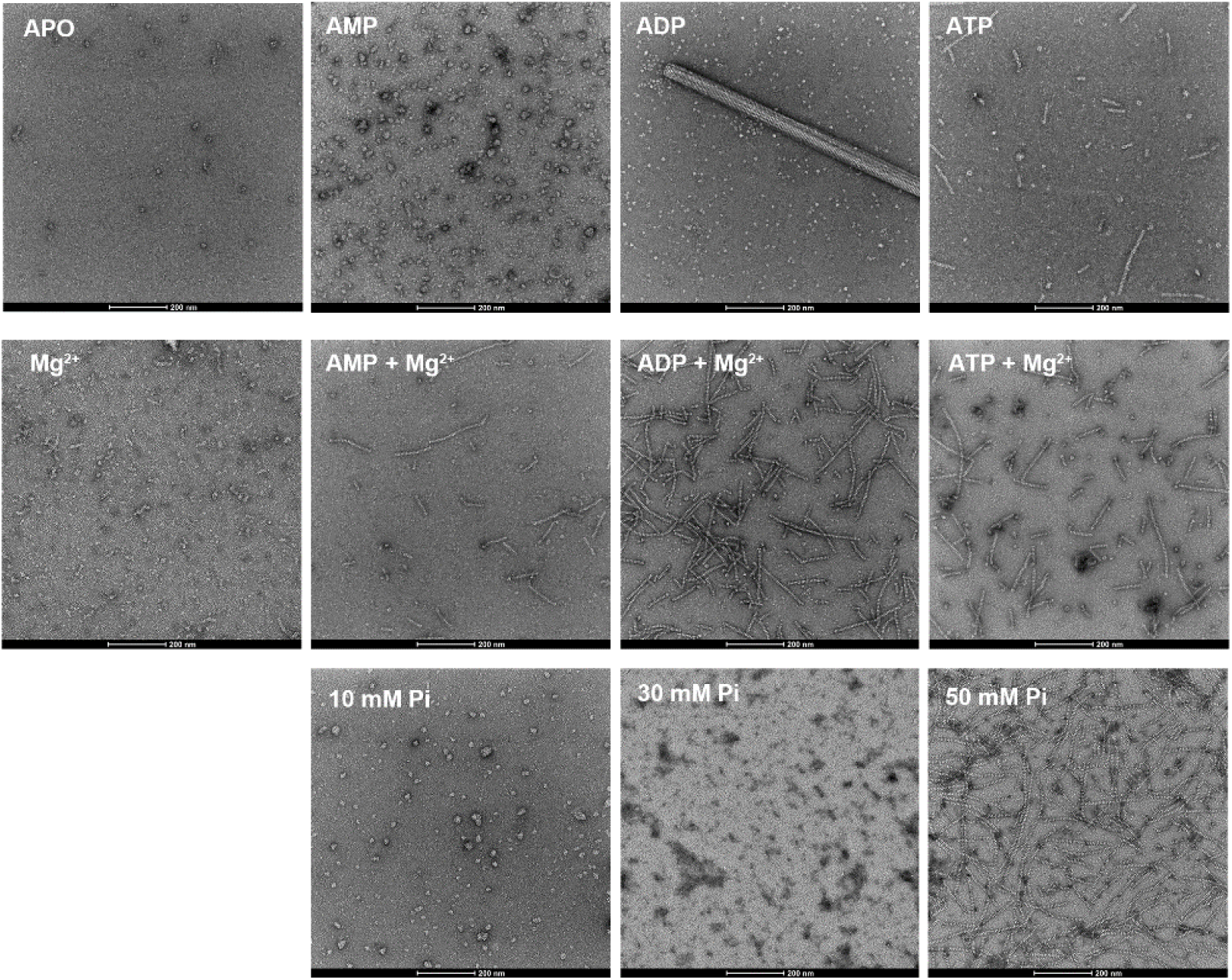
E. coli PRPS assembles filamentous polymers in vitro. Negative staining images of purified E. coli PRPS (1 μM) incubated in various conditions.

**Figure S2.**
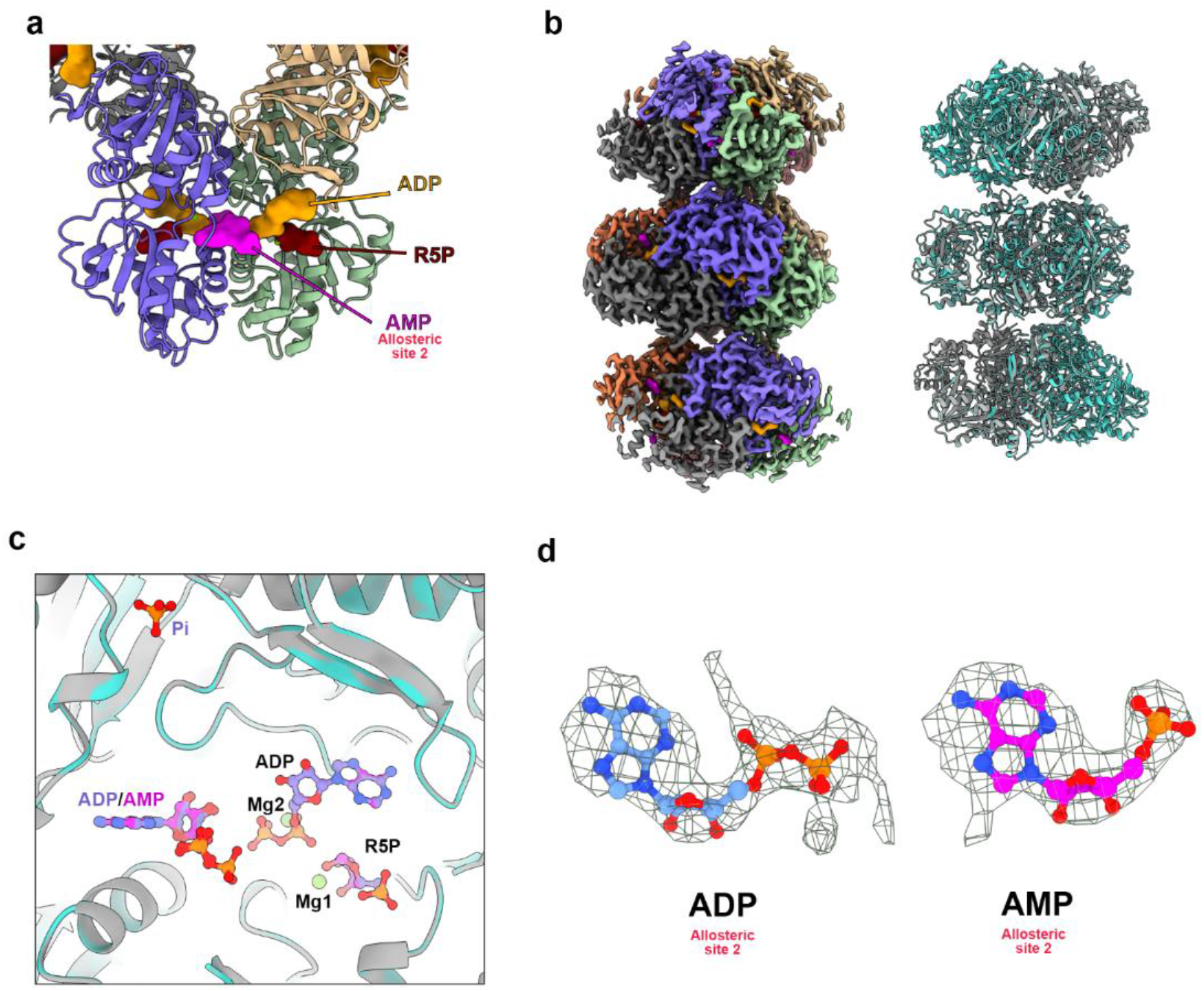
Structure analysis of PRPS type A^AMP/ADP^ filament. (a) The model of the E. coli PRPS parallel dimer in the model of type A^AMP/ADP^ polymer. (b) Cryo-EM reconstruction of type A^AMP/ADP^ (2.6 Å resolution) PRPS polymers (left), overall structure comparison of type A (gray) and type A^AMP/ADP^ (cyan) filaments. (c) Ligand comparison of type A (blue) and type A^AMP/ADP^ (magenta) filaments. (d) The electron density map of ADP (type A) and AMP (type A^AMP/ADP^) in allosteric site 2.

**Figure S3.**
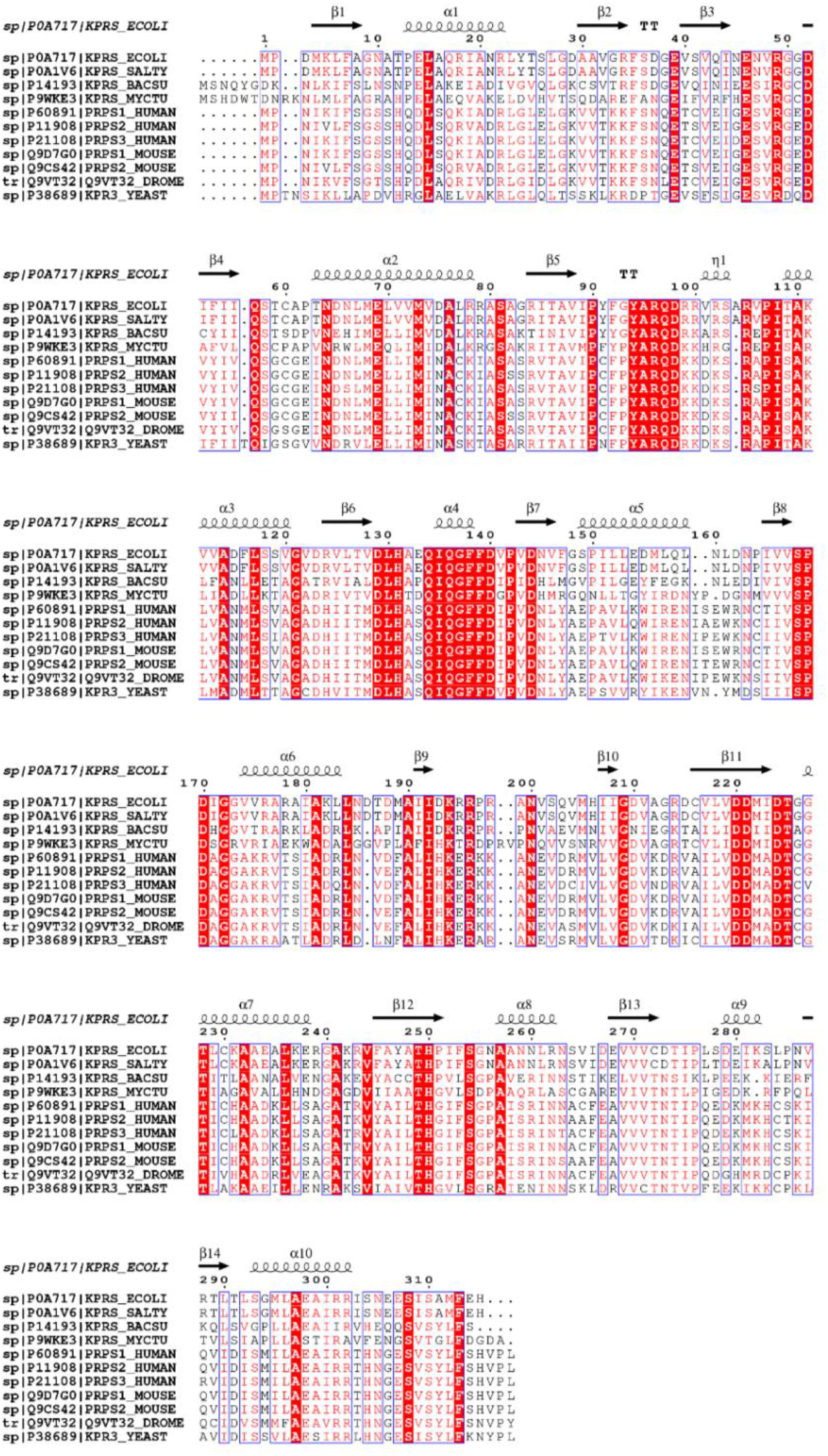
Comparison of PRPS sequences of various organisms.

**Figure S4.**
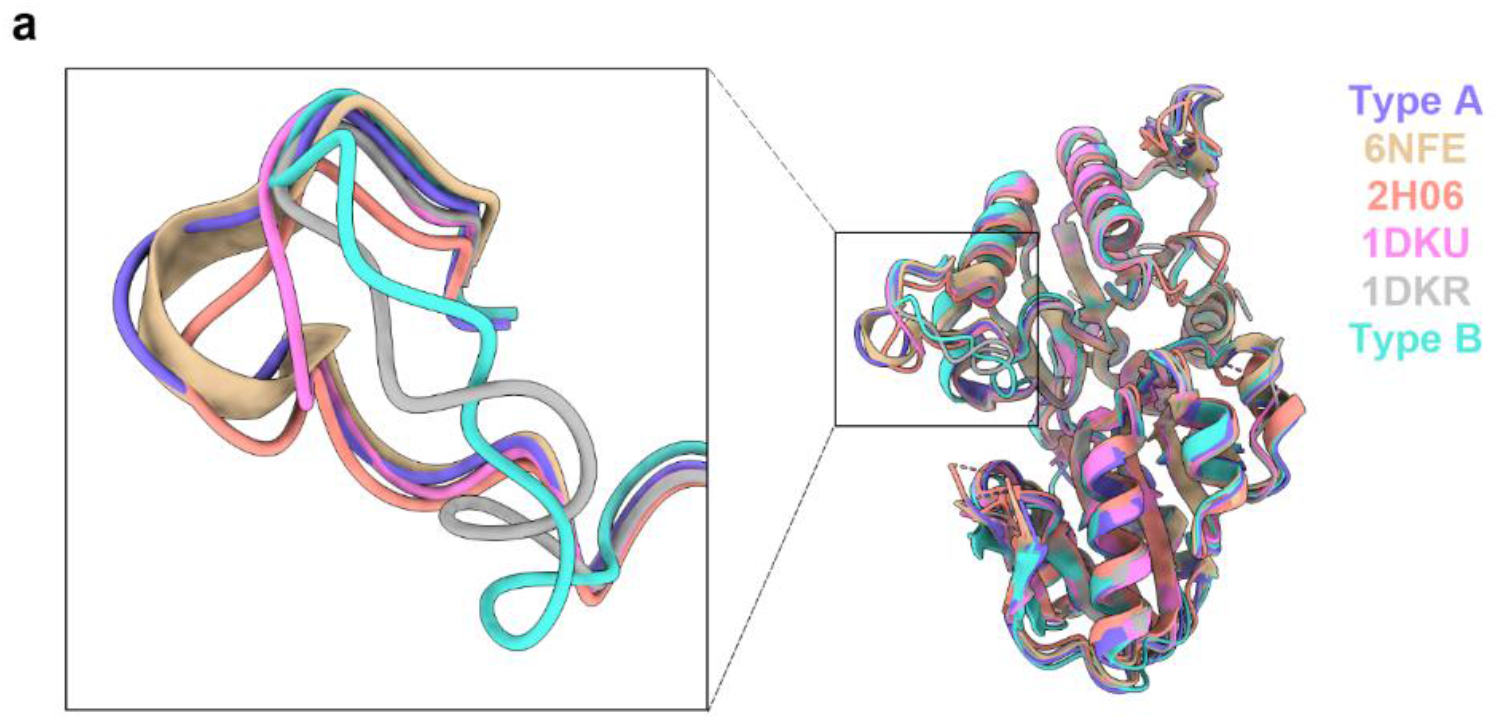
Structure comparison of PRPS in various organisms. The structural comparison of the models of ecPRPS in type A and type B polymers, *Legionella pneumophila* PRPS (6NFE), human PRPS1 (2H06), phosphomethylphosphonic acid adenosyl ester- and methyl phosphonic acid adenosine ester-bound Bacillus subtilis PRPS (1DKU), and Bacillus subtilis PRPS with sulfate ion (1DKR).

**Figure S5.**
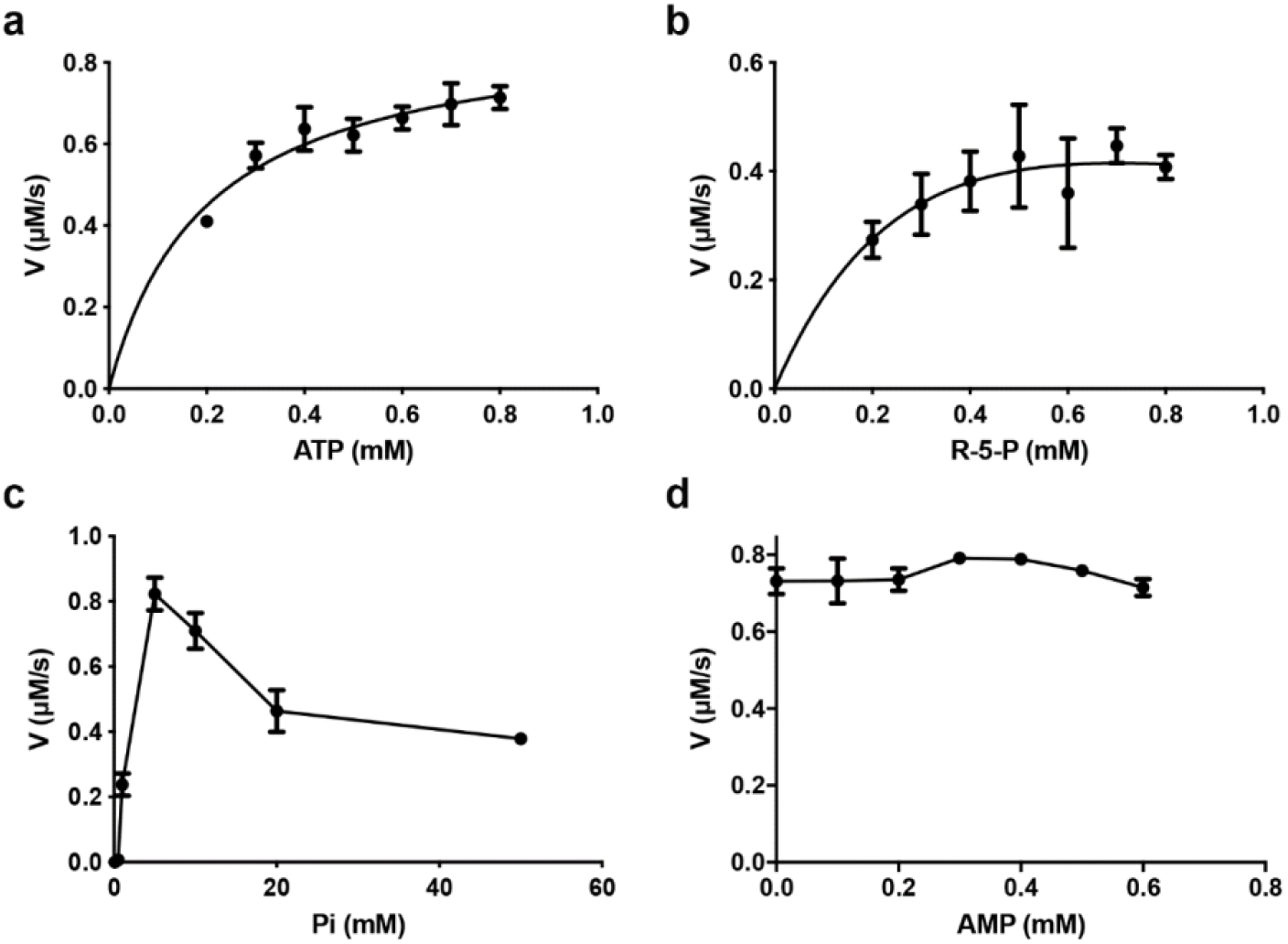
The catalytic activity of E. coli PRPS under conditions with different concentrations of ligands. Graphs show the catalytic activity of wild-type E. coli PRPS in the conditions with various amounts of ATP (a), R5P (b), phosphate ion (c), and AMP (d).

**Figure S6.**
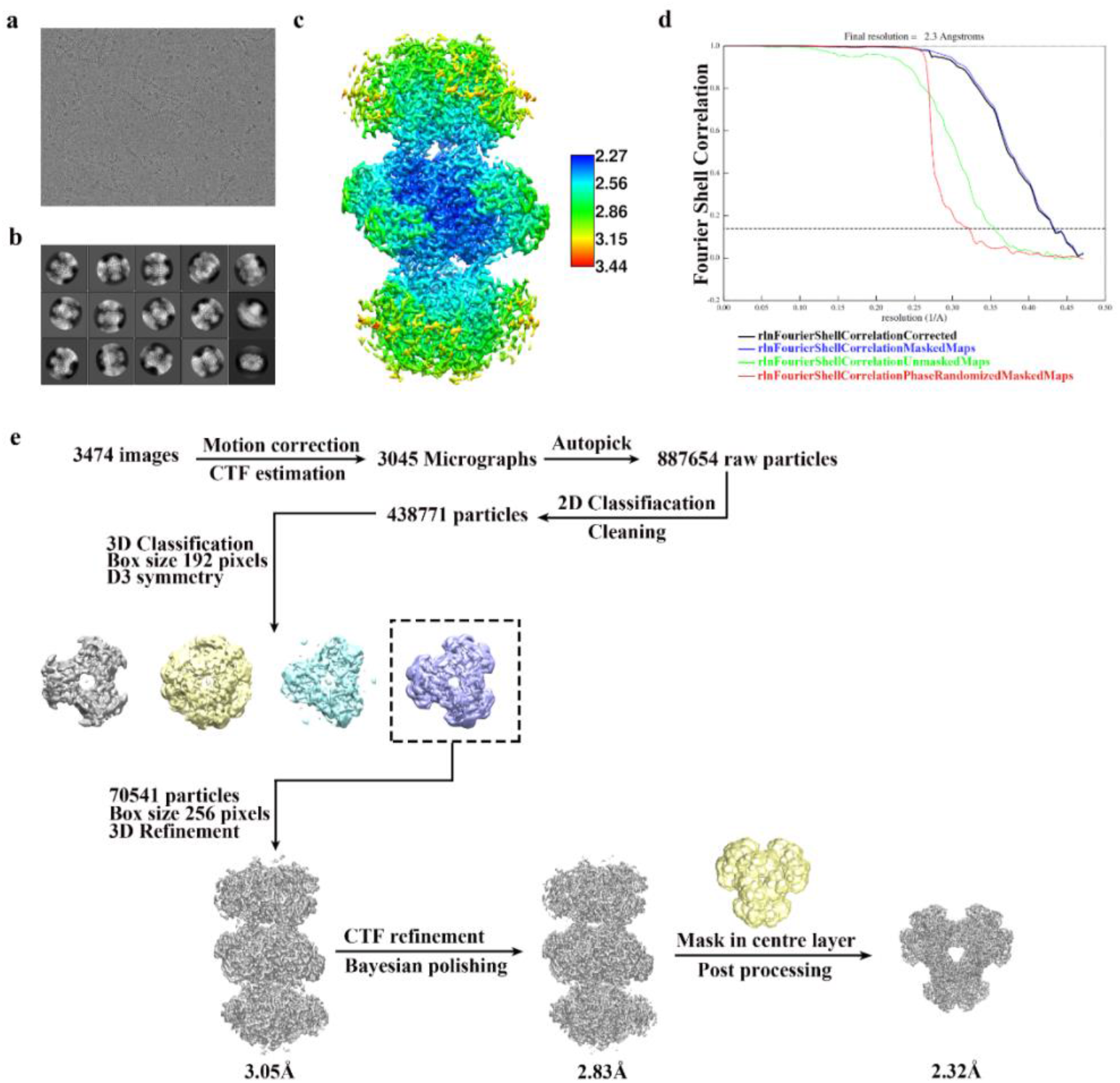
Cryo-EM data processing of PRPS type A filament. (a) Representative Cryo-EM image of PRPS type A filament. (b) Representative 2D averages of PRPS type A filament in different views. (c) Local resolution of the final density map of type A filament. (d) Gold-standard FSC curves of the central hexamer in type A filament density map (dash line shows FSC=0.143). The final average resolution of the hexamer is estimated to be 2.3 Å. (e) Image processing flow chart of PRPS type A filament.

**Figure S7.**
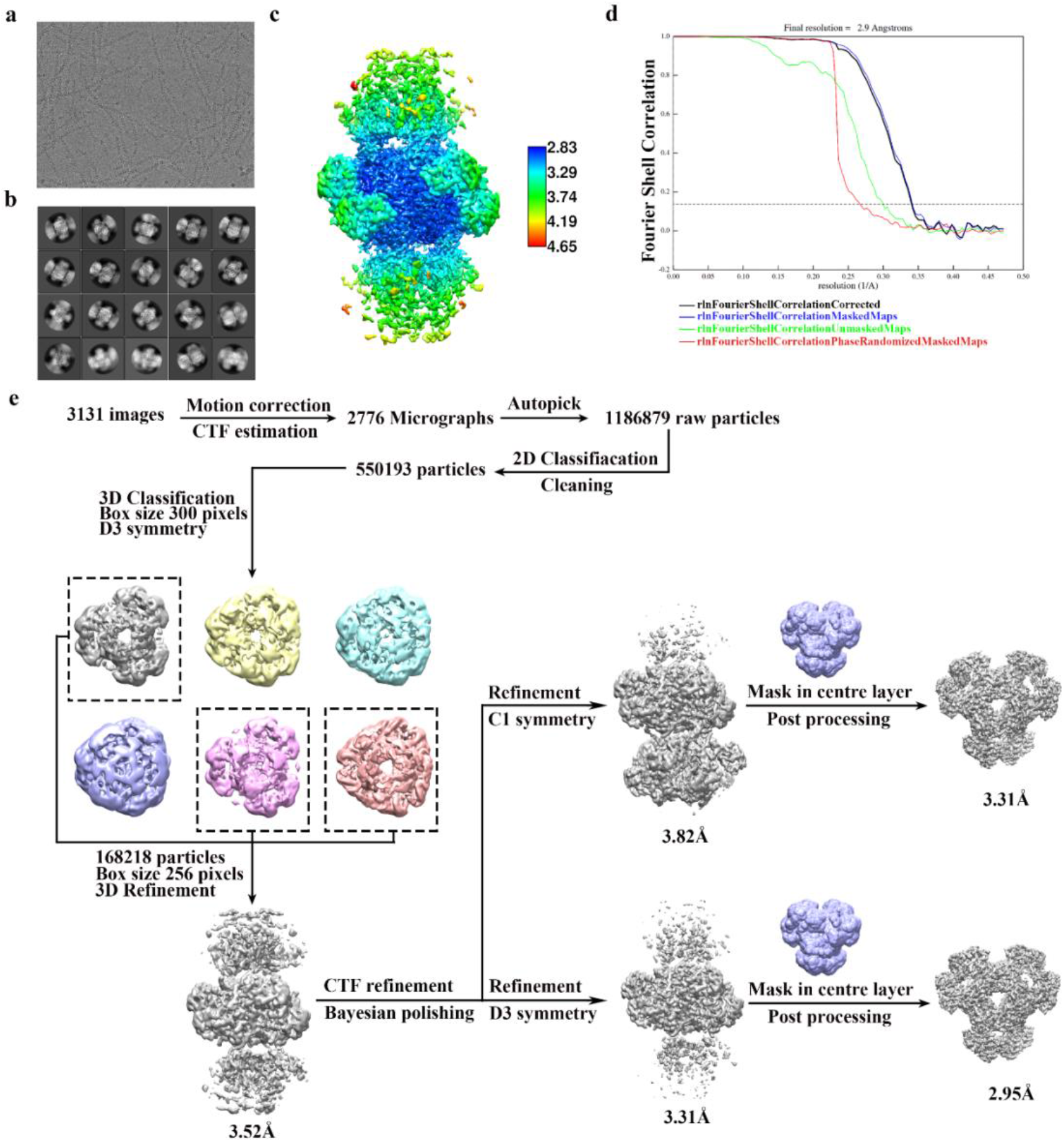
Cryo-EM data processing of PRPS type B filament. (a) Representative Cryo-EM image of PRPS type B filament. (b) Representative 2D averages of PRPS type B filament in different views. (c) Local resolution of the final density map of type B filament. (d) FSC curves of the central hexamer in type B filament density map (dash line shows FSC=0.143). The final average resolution of the hexamer is estimated to be 2.9 Å. (e) Image processing flow chart of PRPS type B filament.

**Figure S8.**
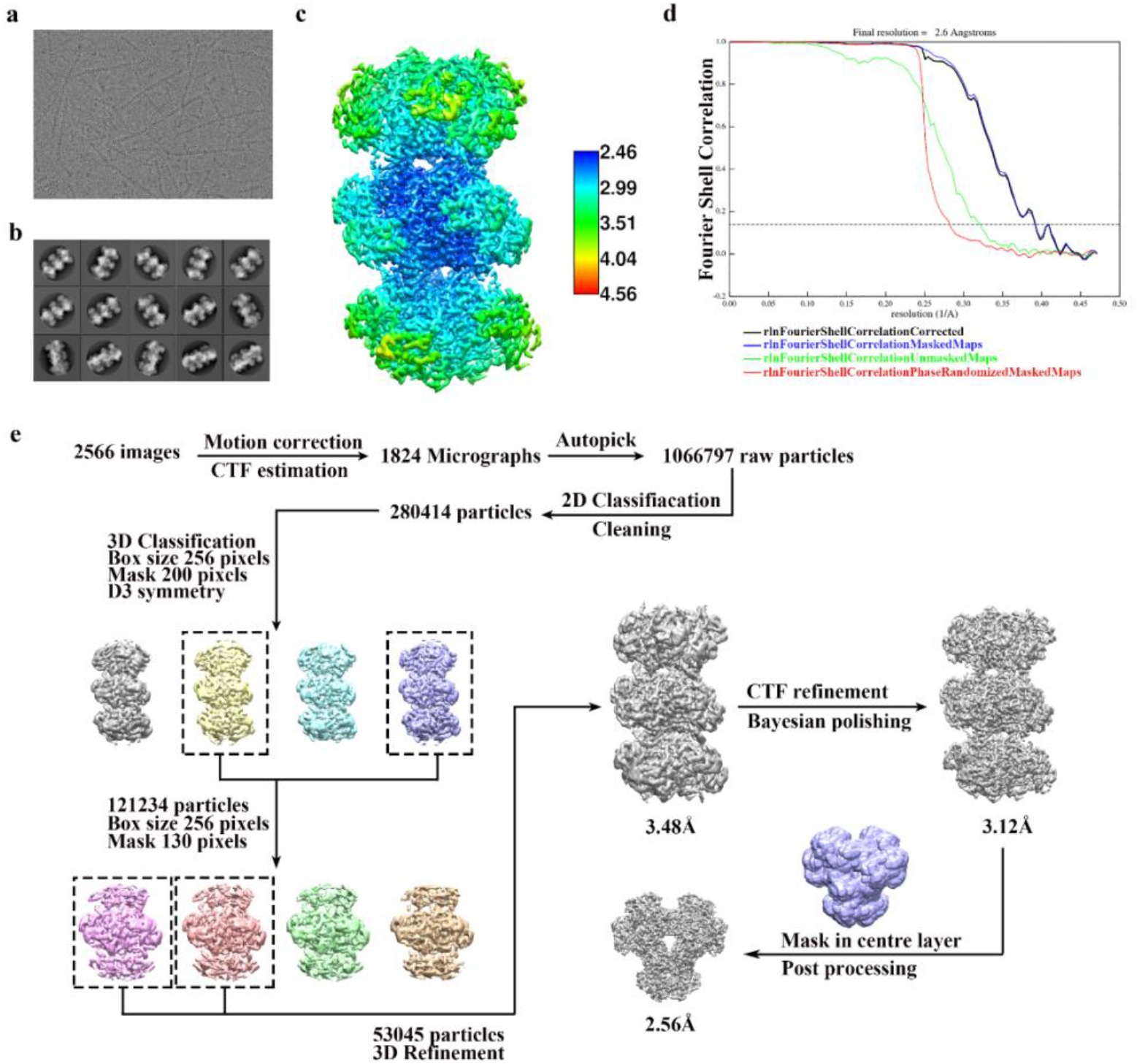
Cryo-EM data processing of PRPS type A^AMP/ADP^ filament. (a) Representative Cryo-EM image of PRPS type B filament. (b) Representative 2D averages of PRPS type A^AMP/ADP^ filament in different views. (c) Local resolution of the final density map of type B filament. (d) FSC curves of the central hexamer in type A^AMP/ADP^ filament density map (dash line shows FSC=0.143). The final average resolution of the hexamer is estimated to be 2.6 Å. (e) Image processing flow chart of PRPS type A^AMP/ADP^ filament.

**Figure S9.**
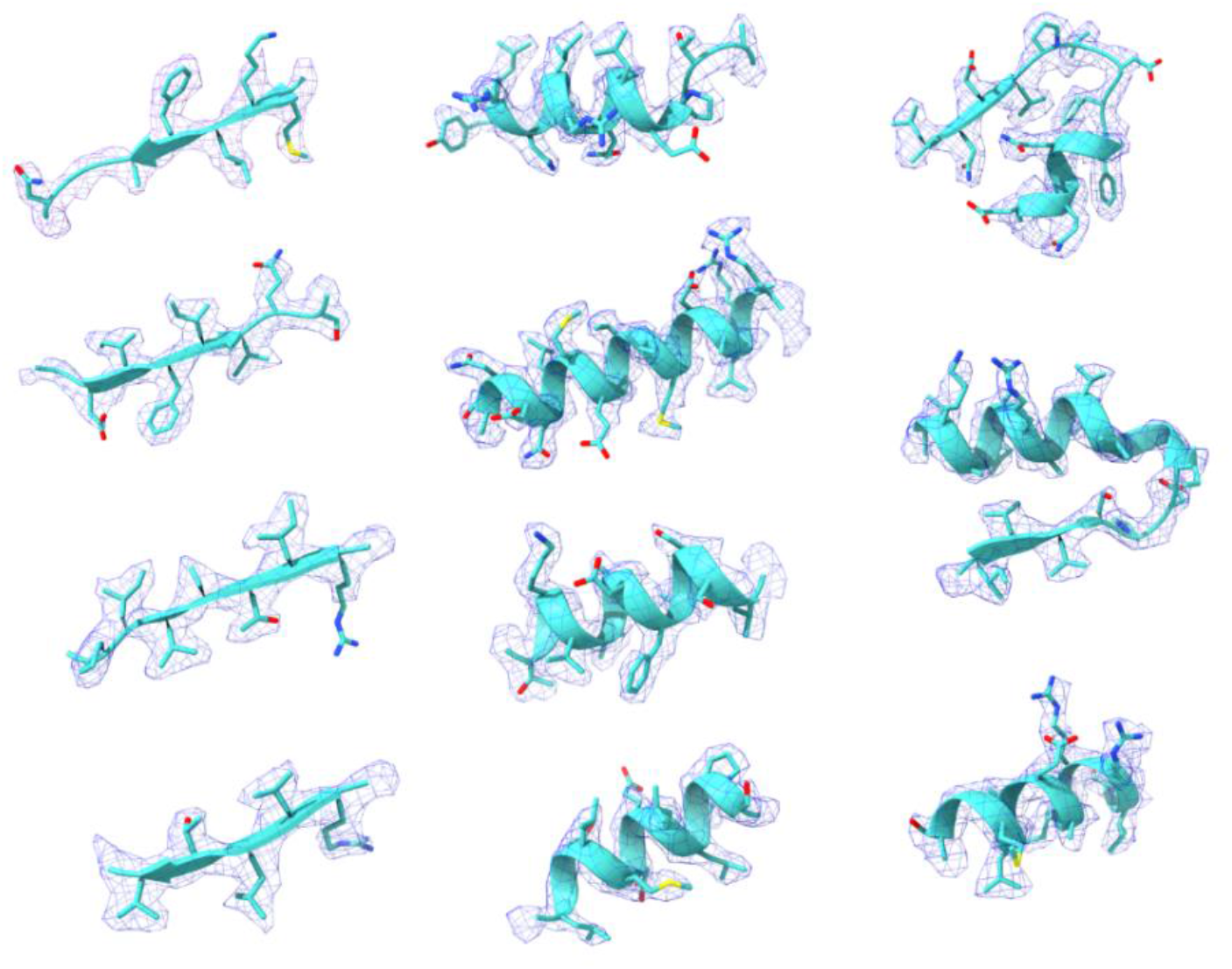
Representative Cryo-EM density map of individual regions of PRPS type A filament.

**Figure S10.**
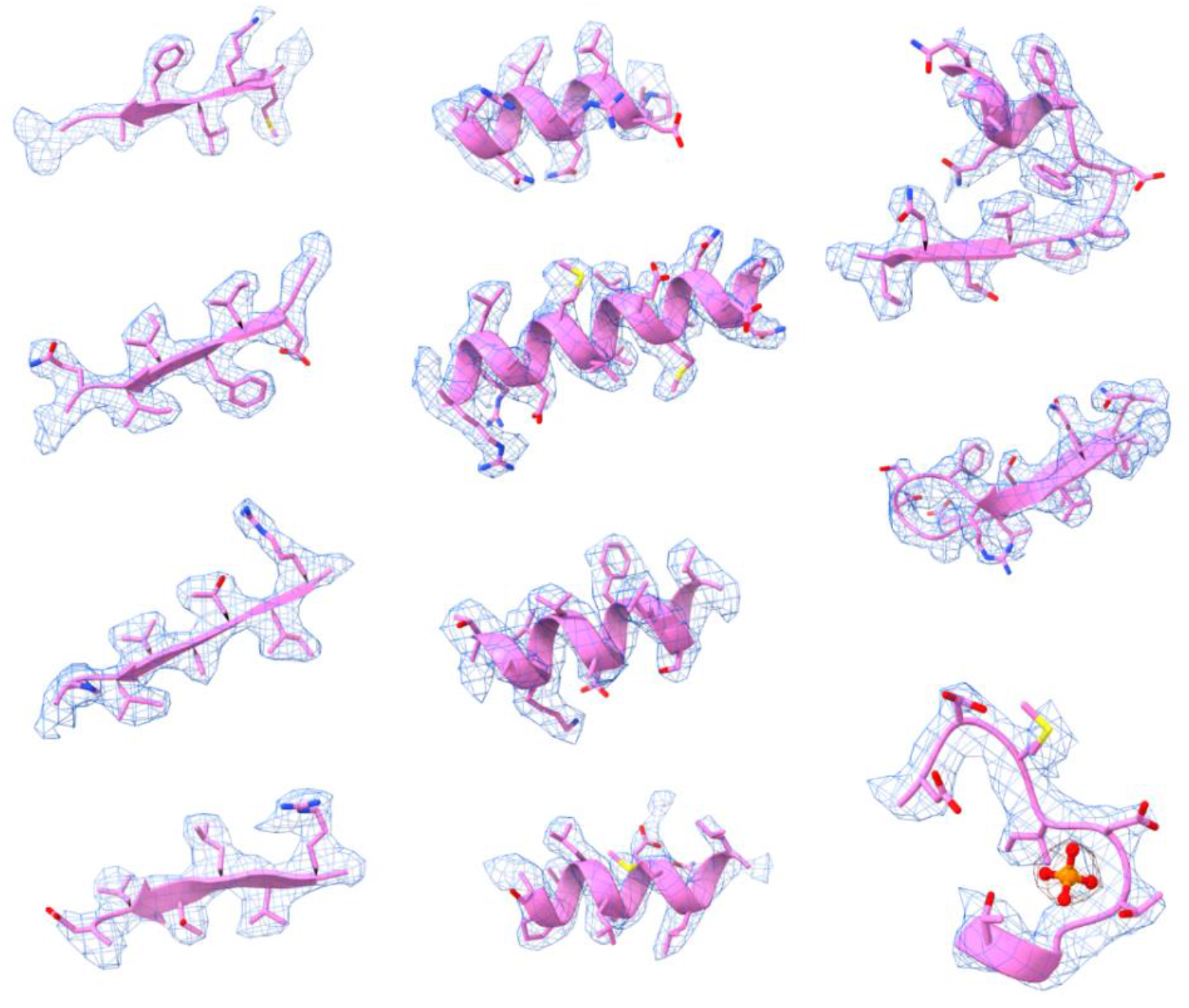
Representative Cryo-EM density map of individual regions of PRPS type B filament.

